# Transduction of mechanical cellular oscillation by the plasma-membrane mechanosensitive channel MSL10

**DOI:** 10.1101/815191

**Authors:** Daniel Tran, Tiffanie Girault, Marjorie Guichard, Sébastien Thomine, Nathalie Leblanc-Fournier, Bruno Moulia, Emmanuel de Langre, Jean-Marc Allain, Jean-Marie Frachisse

## Abstract

Throughout their life, plants are submitted to recurrent cyclic mechanical loading due to wind. The resulting passive oscillation movements of stem and foliage is an important phenomenon for biological and ecological issues such as photosynthesis optimization ^1–3^ and thermal exchange ^4^. The induced motions at plant scale are well described and analyzed, with oscillations at typically 1 to 3 Hz in trees ^5–10^. However, the cellular perception and transduction of such recurring mechanical signals remains an open question. Multimeric protein complexes forming mechanosensitive (MS) channels embedded in the membrane provide an efficient system to rapidly convert mechanical tension into electrical signal ^11^. Here we show that the plasma membrane mechanosensitive channel MscS-LIKE 10 (MSL10) from the model plant Arabidopsis thaliana responds to pulsed membrane stretching with rapid activation and relaxation kinetics in the range of one second. Under sinusoidal membrane stretching MSL10 presents a greater activity than under static stimulation and behaves as a large bandpass oscillation “follower” without filtering the signal in the range of 0.3 to 3 Hz. With a localization in aerial organs naturally submitted to oscillations, our results suggest that the mechanosensitive channel MSL10 represents a molecular component of a universal system of oscillatory perception in plants.

## Main

In animals, transduction of vibrational stimulation is achieved through MS channels in organs with specialized structures, such as the ear in which the different frequencies are spatially separated^12^, or by the organ motion as in touch sensation^13^. In plants, such specialized features have not yet been reported, and it remains unclear whether and how MS channels participate in the perception of oscillatory stimuli. To investigate this question, we studied MSL family members, homologues of the Mechanosensitive channel of Small conductance (MscS) from *E. coli* ^14,15^, as they are found in all land plant genomes ^16^. We have focused our study on MSL10, the most widely expressed, plasma membrane-localized and functionally characterized member in Arabidopsis thaliana ^17,18^.

To characterize the response of Arabidopsis to wind mechanical stimulation, we examined the frequency of free oscillations of plant with young flowering stem subjected to a short air pulse (supplementary movie 1). Using the Vibration Phenotyping System (https://vimeo.com/213665517)^19^, we determined the image correlation coefficient depicting the pendulum movement of the stem on 6 plants and obtained a mean frequency of 2.8±1.0 Hz (mean ±SD, n=131) (Fig. 1a). This frequency is inside the one excited by the wind^20^. Then, to determine whether MSL10 localization is compatible with a function as oscillation sensor, we characterized its expression pattern on plants at a flowering stage. GUS reporter driven by *MSL10* promoter was present in steam and leaf vasculature and at the root tip (Fig. 1b, c) ^17^. This expression pattern of *MSL10*, especially at the junction between roots and shoots, which experience the major tension induced by leaves and stem motion, is an expected location for probing the motion induced by the wind (Fig. 1d).

**Figure 1.**
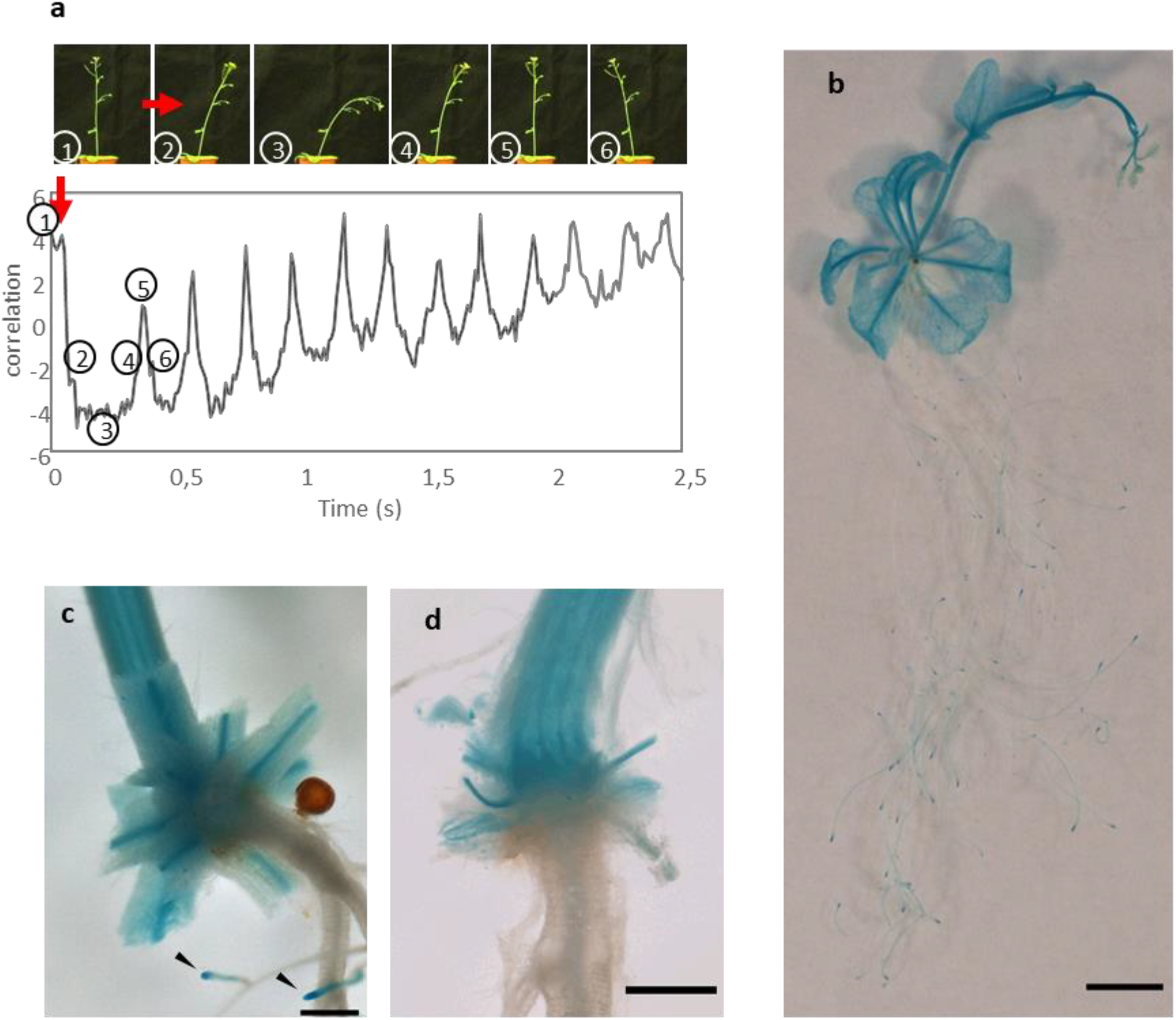
Oscillatory movement and *MSL10* expression pattern in aerial part of Arabidopsis plants. **a,** Images of the oscillatory movement of the stem induced by an air pulse of 60 ms, the correlation coefficient curve visualize the oscillating and the damping of the stem movement (red arrow: air pulse), **b-d** Blue staining represents the β-glucuronidase (GUS) activity driven by the promoter of *MSL10*. The *MSL10* promoter drives expression of the reporter gene **a,** in the root tip (indicated by arrows in c) and throughout vasculature of the leaves and stem, **c,** at the bottom of leaf petioles and **d,** in the root-stem junction (d is the same view as c, with petioles entirely removed). Scale: b, 5mm; c and d, 500μm.

Channel kinetic properties are crucial for its ability to perceive oscillatory stimulation at various frequencies. In order to know how fast MSL10 responds to rapid variations in membrane mechanical tension, we characterized the kinetics of this channel using the patch-clamp technique. To specifically monitor MSL10 activity in its endogenous environment, we expressed the *MSL10* gene in protoplasts from a quintuple mutant (*Δ5*) lacking the activity of five *MSL*-encoding genes (*msl4;msl5;msl6;msl9;msl10)*. This provides a low background to record mechanically-activated currents from *MSL10*-expressing protoplasts (*Δ5*+*MSL10*) ^17^ by applying pulses of pressure whilst monitoring transmembrane currents at a constant voltage (−186mV) on a membrane excised patch in outside-out configuration (Fig. 2a). At this physiologically relevant membrane potential, opening of a single stretch-activated channel caused a current variation of 19.4 ± 1.7 pA (Fig. 2a, n=14) as reported in root protoplasts expressing MSL10^17,21^. The sustained activity under the membrane tension of MSL10 without inactivation clearly distinguishes it from the plasma membrane rapidly activated calcium MS channel activity (RMA), which displays rapid inactivation ^22,23^. We observed that the activation of MSL10 current increased exponentially in response to pressure, with time constants τ_act_ ranging from 1000 ms at 30 mmHg to 200 ms at 100 mmHg (Fig. 2b and Supplementary Fig. 1, n = 15). The current-pressure relationship representing the MSL10 channel sensitivity to membrane tension was well described by a Boltzmann function with a pressure for half-activation (P_0.5_) of 49.3 ± 3.4 mmHg, an activation threshold of about 30 mmHg and a saturation pressure of about 70 mmHg (Fig.2c and Supplementary Fig. 2a-c).

**Figure 2.**
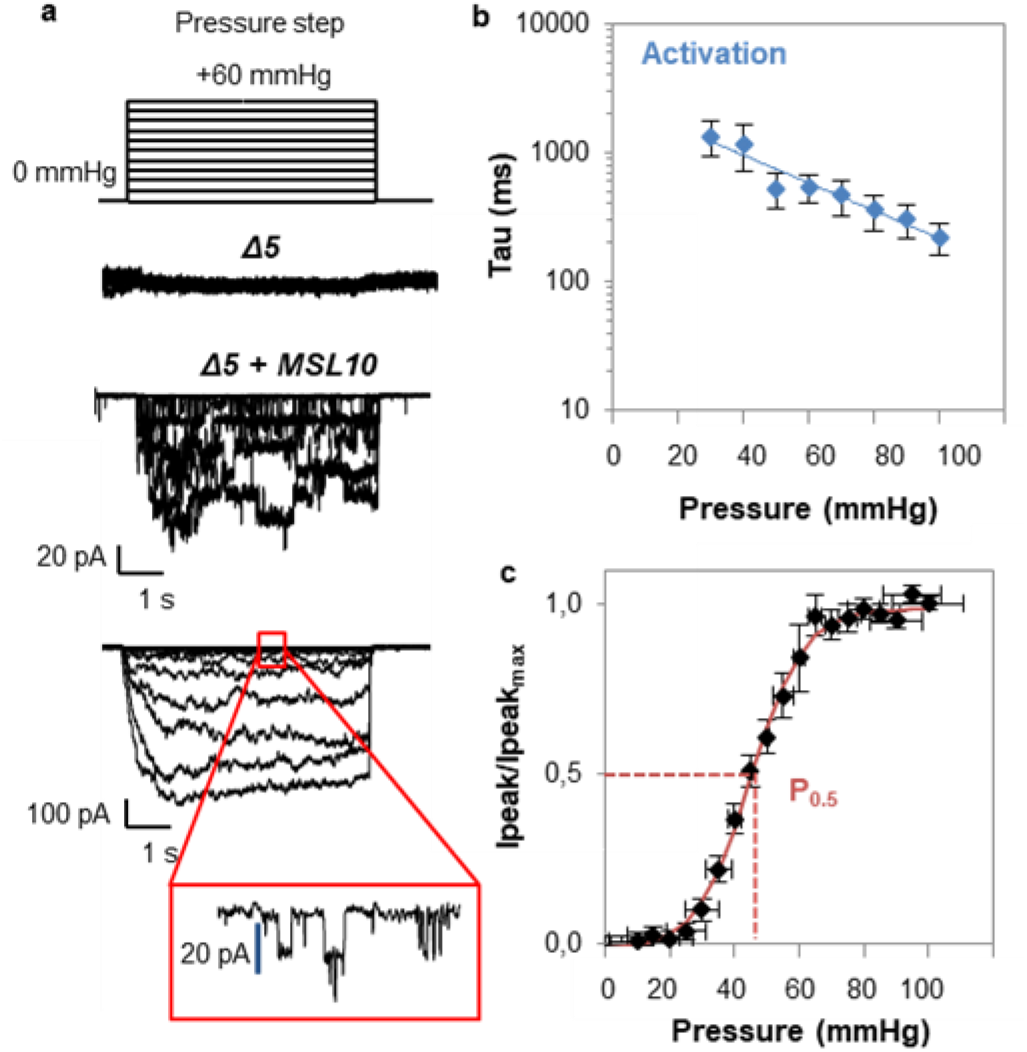
Gating kinetics and pressure dependence of MSL10 in native membrane. **a**, Quintuple mutant **(Δ5)** stimulated by increasing pressure steps in outside-out patch configuration shows no mechanically activated current. Δ*5* mutant expressing MSL10 (Δ*5+MSL10*) shows currents stimulated in outside-out configuration by increasing pressure steps with slow activation kinetics. The current amplitude varies from one patch to another depending on channel density (expression of the protein). Single channel amplitude shows current transitions of 19.4 ± 1.7 pA at −186 mV (n = 14 protoplasts). **b**, Pressure dependence of activation time constant for MSL10 channel in excised outside-out-patch configuration (see Extended Data Fig. 1). Results are normalized with the P_0.5_ of each patch and data represent mean ± S.E.M (n = 15 protoplasts), **c, I_max_** normalized current-pressure relationship of stretch-activated currents in excised outside-out-patch configuration in Δ*5+MSL10*, fitted with a Boltzmann equation. P_0.5_ of 49.3 ±3.4 mmHg is the average value determined for individual cells. Data represent mean ± S.E.M (n = 15 protoplasts). The membrane potential is clamped at −186 mV. MSL10 protein is transiently expressed in quintuple *msl4;msl5;msl6;ms19;msll0* mutant *(Δ5)* protoplasts. Ionic conditions are described in the Materials and Methods.

We then examined the effect of oscillatory membrane tension on MSL10 channel activity. To do so, MSL10 activity was recorded under oscillating pressures at a wide range of frequencies from 0.3 to 30 Hz (supplementary movie 2, Fig. 3a). Whatever the frequency tested, opening events occurred almost exclusively during the upper phase of the period (≥ 80 % of cases) (Fig. 3b, Supplementary Fig. 3). At low frequency (≤ 1 Hz), at least one opening transition of the channel was triggered during each period, (Fig. 3c, 100 % of cases), at 3 Hz 70 % of the periods triggered channel opening, while at 30 Hz only 20 % of the periods were efficient (Fig 3c and Supplementary Fig. 3). Thus, the channel does not open randomly in response to oscillatory stimulations, even when some pressure pulses at 3 Hz and beyond were not efficient to trigger the channel opening (Supplementary Fig. 4).

**Figure 3.**
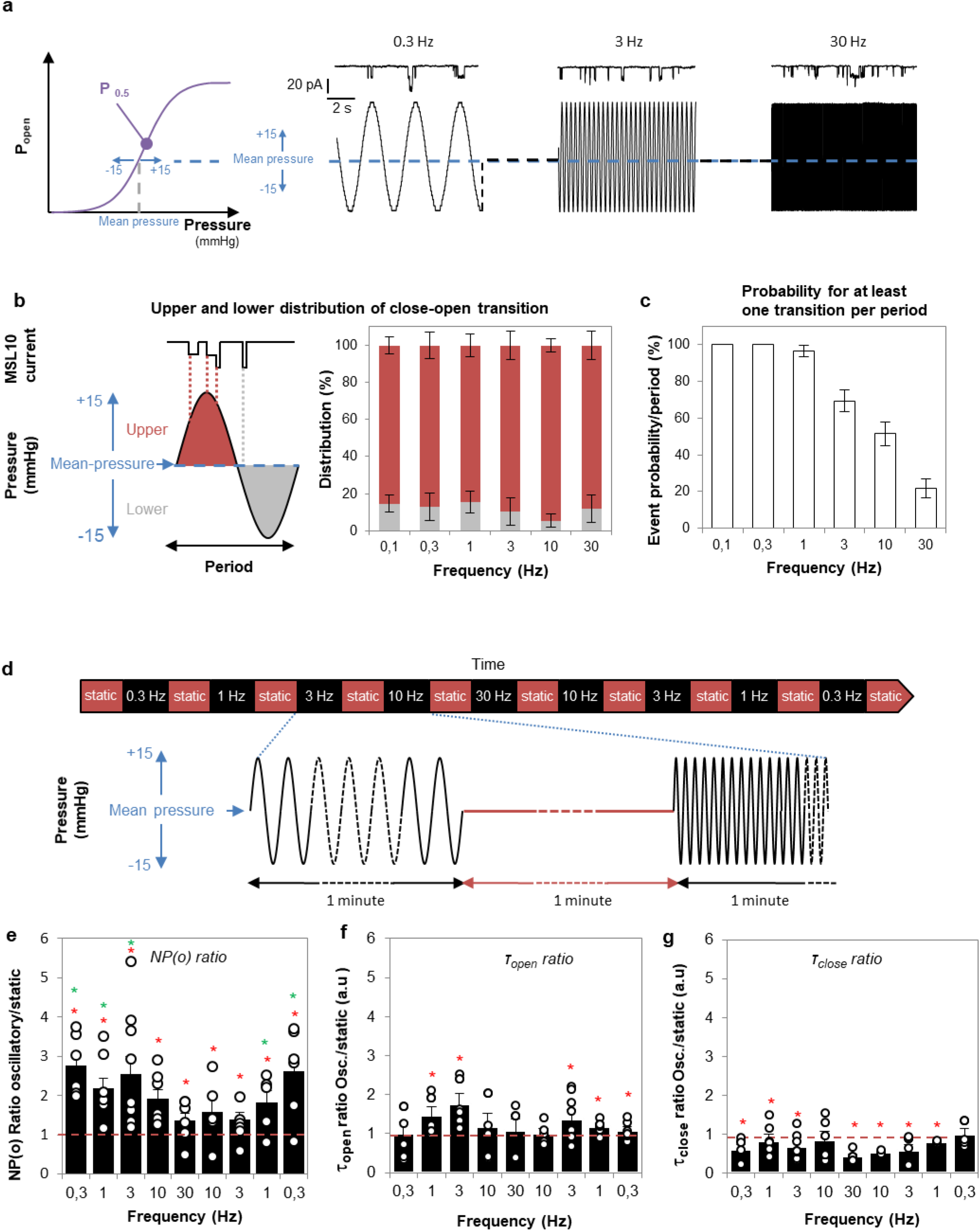
Effect of oscillatory pressure stimulation on MSL10 channel characteristics. **a**, Representative recording of single channel activity of MSL10 in excised outside-out-patch configuration, in response to oscillatory pressure stimulation at 0.3 Hz, 3 Hz and 30 Hz. An oscillatory pressure of +15/-15 mmHg from a mean pressure is applied. b, *Left*, Idealized MSL10 current used for event analysis in response to oscillatory pressure. *Right*, Distribution of close-open transitions (at least one) elicited at upper (red bars) or lower (light grey bars) pressure as a function of frequency for 15 seconds of loading. Results represent mean ± S.E.M (n ≥ 5 protoplasts). c, Probability that MSL10 channel undergoes at least one close-open transition per period as a function of frequency. Results represent mean ± S.E.M (n ≥ 5 protoplasts). d, Sequences of 1 min oscillatory pressure alternating with 1 min static pressure are performed on excised outside-out-patches over time. The oscillatory stimulation (same protocol for Supplementary Fig. 6a-c) is of 30 mmHg amplitude (+15/-15 inmHg from a mean-pressure level) with a sweep of frequencies from 0.3 to 30 Hz (--), while static stimulation is at mean-pressure (--). **e-g**, Relative effect of frequency (oscillatory/static mean-pressure) on **e**, open probability NP(o), **f**, open state time constant and **g**, closed state time constant. The red dashed line represents the relative ratio static/static (=1). Each point represents each biological replicate (n ≥ 5 for a given frequency); asterisk in red (*) indicates in **e, f** and **g** that mean value is significantly different from **1** (Mann-Whitney Rank Sum Test, p<0.05), asterisk in green (*) in **e** indicates mean NP(o) is significantly different from mean NP(o) obtained at 30 Hz (Mann-Whitney Rank Sum Test, p<0.05). We have determined *NP(o)_osc_* for each oscillatory sinusoidal pressure frequency (0.3 to 30 Hz) and *NP(o)_stat_* for each static stimulation prior to frequency stimulation. In 3b we present the ratio *NP(o)_osc_/NP(o)_stat_* called *NP(o) Ratio*. The same principle is applied for *T_open_ ratio* and *T_c]ose_ ratio* in 3c and d. Other conditions same as Fig. 2.

We then undertook a comparison between static and oscillatory stimuli for a same applied “mean-pressure” using a protocol alternating the two types of stimulations (see Fig. 3d). A static stimulation held at “mean-pressure” was applied for 1 min followed by a sinusoidal pressure stimulation of +15/−15 mmHg from the mean-pressure baseline at a given frequency for 1 min (Fig. 3d). This protocol was repeated to sweep frequencies from 0.3 to 30 Hz and then from 30 to 0.3 Hz, always with the same mean-pressure, in order to determine the effect of frequencies on channel activity compared to that under prior static stimulation (Fig. 3d). Figure 3e-g shows the relative effect of frequencies (ratios oscillatory/static) on channels NP(o), τ_open_ and τ_close_ on at least 5 membrane patches.

A ratio (NP(o) osc/ NP(o) stat) above 1 indicates a greater activity of the channel under sinusoidal stimulation than under static stimulation. We observed that at each frequency, the NP(o) ratio is significantly greater than one, meaning that the mean open probability is significantly higher upon dynamic than static stimulation, while the pressure applied was on average the same (Fig. 3e; red asterisks, Mann-Whitney Rank Sum Test, p≤0.05). The highest ratios are observed at low frequency (0.3, 1 and 3 Hz) corresponding to the frequencies of plant oscillation measured in Figure 1a (Fig. 3e, green asterisks, Mann-Whitney Rank Sum Test, p≤0.05). The asymmetry observed in NP(o) distribution for decreasing and increasing frequencies (Fig. 3e and supplementary Fig.5a) likely reflect the diminution of the active channels over time of the experiment. Supplementary Figure 5 exemplifies the effect of frequencies on NP(o), τ_open_ and τ_close_ obtained for a representative recording. Under oscillatory stimulation NP(o) increased, τ_open_ were unchanged while τ_close_ were decreased compared to the static stimulation. In order to further quantify the opening and closing oscillation dependency of MSL10, we compared open and close time constants obtained on five patches, either under static or dynamic conditions. We measured a mean open time constant in static condition of τ_open_ static = 14.7 ± 1.9 ms (n ≥ 5). This time constant is not or weakly affected by oscillatory stimulation with a τ_open_ oscillation relative to static above 1 (Fig. 3f). The mean close time constant τ_close_ decreased significantly from τ_close_ static = 164.5 ± 24.8 ms to τ_close_ oscillation = 106.4 ± 17.6 ms (all frequencies, n ≥ 5). This is reflected by the ratios τ_close_osc/τ_close_ stat lower than 1 (Fig. 3g), pointing to the fact that MSL10 spends less time in the closed state due to an increase in their opening probability upon oscillatory stimuli.

Mammalian Piezo1 and Piezo2 have been reported as pronounced frequency filters^13^, thus allowing transduction of repetitive mechanical stimuli at a given frequency. This was attributed to their strong inactivation. In MSL10, we didn’t observe a strong inactivation, but still we observed a clear oscillation dependence in a wide range of frequencies. We tested if this may come from the channel natural kinematic of opening and closing as a function of the tension. To do so, we implemented a two-states model (see Material & Methods), which fits well our data despite its lack of explicit frequency dependency (Fig. 4a). We observe an oscillatory dependence of the response, as the model predicts a higher NP(o) ratio between oscillatory and static pressure at low frequency than at high frequency.

**Figure 4.**
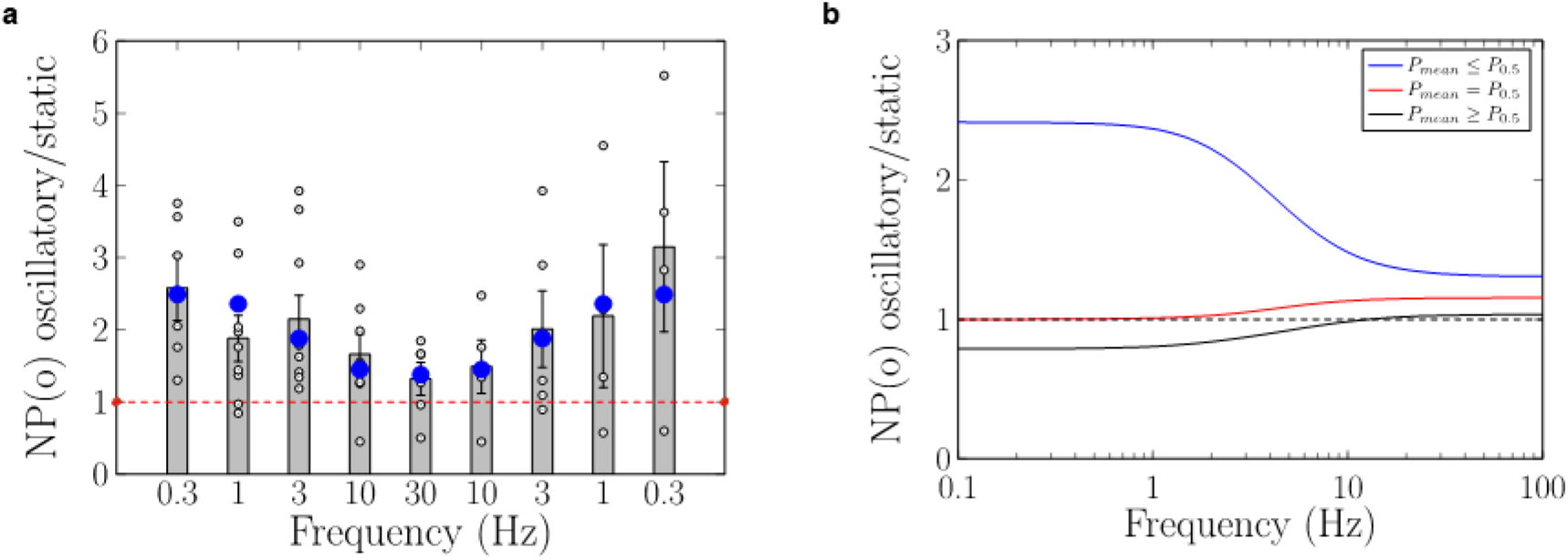
Modelling of MSL 10 channel as a classical double state system. The channel is modelled as an 2 states (open-close) system, with rates classically changing exponentially with the applied pressure. No specific frequency dependence is introduced. **(a)** Adjustment of the model to experimental data. Model predictions (blue circles) are superimposed over data from Fig. 4e. **(b)** Prediction of the NP(o) ratio between oscillatory/static mean-pressure for different initial mean-pressure, using the same parameters. Blue: parameters obtained from the experiment (mean-pressure 7.6mmHg below P_0.5_). Red: mean-pressure taken as P_0.5_. Black: mean-pressure increased by 7.6mmHg with respect to P_0.5_.

The low frequency response of the channel can be explained physically by the non-linear response of the channel to static pressures (the Boltzmann curve). As the mean-pressure is below P_0.5_, an increase in pressure will open more channels than the same decrease in pressure. Thus, it is expected that at low frequencies the ratio is greater than one. Choosing an initial mean-pressure exactly at P_0.5_ will have symmetrized the effects of the increase and decrease in pressure leading to a ratio of 1 (see Fig. 4b). Similarly, starting from a pressure above P_0.5_ will have led to a decrease of the ratio. The reliability of this model raises two questions: at which frequencies the channel is solicited in vivo and what is the mean pressure applied to the membrane in vivo? The free oscillations of Arabidopsis around 3 Hz that we have measured (Fig. 1a) are within the range of low frequencies presented in the model for which oscillatory stimulations are more efficient than static one (Fig. 4b), and are well described by the Bolzman response. For the mean pressure applied in vivo, one should expect to have two contributions: a baseline due to the turgor pressure, and one due to the mean bending during the plant oscillation, proportional to the amplitude of the plant oscillations. Thus, for a plant in rest condition, not submitted to wind loading, the MSL10 channel is expected to be in the very low domain of solicitation of the Boltzmann curve (Fig. 2c) for which the channel is closed. At low amplitudes of oscillation, corresponding to a solicitation in the domain of the Boltzmann below P_0.5_, the channel will be more active under movement than during an equivalent static bending and thus will transduce oscillation into cellular ion fluxes. At high amplitude of oscillation, corresponding to a domain above P_0.5_ on the Boltzmann curve, the channel will be less active during oscillation than during a static bending. This might represent a homeostatic behavior amplifying channel activity at low amplitude but decreasing channel activity at large amplitude.

At high frequency, we predict ratio larger than 1, whatever the initial mean-pressure. This effect is due to higher pressure sensitivity for opening than for closing the channel, but is harder to explain intuitively. Interestingly, the characteristic frequency, at which the channel changes from one regime to the other one, doesn’t seem to depend on the mean-pressure. This effect on the channel observed for mechanical stimulation at frequency higher than 10 Hz is difficult to rely to a cellular physiological function.

Our finding also raises the question on how oscillations occurring at the scale of the plant organ could be relayed at the scale of the cell membrane. We know that a mechanical stimulation, in order to be efficient (in term of physiological response), should produce a tissue/cell deformation^24,25^. In previous study on Arabidopsis^26^, sinusoidal sweep excitation, mimicking wind, combined with high-speed imaging allowed us to estimate several modal frequencies and the corresponding spatial localizations of deformation. The spatial localizations of the deformation are compatible with the localizations in the plant of MSL10 as measured here (Fig. 1b). Therefore, to link membrane and organ scales we propose a qualitative model in which tissue/cell deformation induced by mechanical oscillations would induce local membrane tension able to trigger MSL10 channel. However, a full assessment of this hypothesis requires working out the full “localization/distribution/intensity” of the membrane stretching or tensions.

## Conclusion

In plants, the functions of plasma membrane-located MSLs are unknown, with the exception of MSL8 which was shown to be involved in pollen hydration^27^. This is particularly surprising for MSL10 as it is the most studied member of the MSL family. Actually, MSL10 was shown to induce cell death, but this effect was found to be separable from its mechanosensitive ion channel activity^28^. In the present study we provide compelling evidences supporting that MSL10 acts not only as a classical transducer of sustained force but also as a transducer able to translate mechanical oscillations. With a selectivity in favor of anions^17,18^ the stretch-activated channel MSL10 is a potent actor of the plasma membrane depolarization. Thus, solicitation of MSL10 via mechanical stress delivered as sustained or even more efficiently as repetitive load to the membrane is a favorable situation to initiate electrical signaling.

This study supports that MSL10 might represent a molecular component of a system of oscillatory perception in plants. Our findings open new avenues for studying the molecular mechanisms involved in perception of oscillations that allows environmental adaptation.

## Methods

### Histology

Transgenic *Arabidopsis* lines used for histochemical studies and carrying the *pMSL10*::*GUS* promoter-reporter gene fusion were obtained previously^17^. In order to perform detection of β-glucuronidase (GUS) activity on whole plant, plants were grown on agar plate. In this culture condition, mature plants with flowering stem have a reduced height of ~4 cm and are suitable for staining. Tissue was fixed for 30 min in ice-cold 90% acetone, then incubated overnight at 37°C in 0.5 μg/mL 5-bromo-4-chloro-3-indoyl β-glucuronic acid, 100mM NaPO_4_ (pH 7), 0.1% Triton X-100, 5 mM potassium ferrycyanide, and 10mM EDTA. Samples were then dehydrated through an ethanol series and photographed with a camera or with a Nikon AZ100 MultiZoom macroscope (objective: AZ-Plan Apo 1X NA 0.1 WD 35 mm (Nikon)).

### Callus initiation and maintenance

*Arabidopsis thaliana* (Col-0 accession) surface-sterilized seeds were sown on “initiation medium” containing 4.3 g/L Murashige and Skoog salts (MS, Sigma-Aldrich), 2% sucrose, 10 mg/L myo-inositol, 100 μg/L nicotinic acid, 1 mg/L thiamine-HCl, 100 μg/L pyridoxine-HCl, 400 μg/L glycine, 0.23 μM kinetin, 4.5 μM 2,4-D, 1% Phytagel (pH 5.7). For callus generation, seeds were cultured in a growth chamber for 15 days. Calli were then transferred onto “maintenance medium” containing 4.3 g/L MS salts (Sigma-Aldrich), 2% sucrose, 10 mg/L myo-inositol, 100 μg/L nicotinic acid, 1 mg/L thiamine-HCl, 100 μg/L pyridoxine-HCl, 400 μg/L glycine, 0.46 μM kinetin, 2.25 μM 2,4-D, 1% phytagel, (pH 5.7), and sub-cultured every 15 days onto fresh “maintenance medium”.

### Protoplast isolation and transient transformation

Calli from *Arabidopsis* were digested for 15 min at 22 °C under hyperosmotic conditions (2 mM CaCl_2_, 2 mM MgCl_2_, 1 mM KCl, 10 mM MESs (pH 5.5), 0.2 % cellulysine (Calbochem), 0.2 % cellulase RS (Onozuka RS, Yakult Honsha Co.), 0.004 % pectolyase Y23 (Kikkoman Corporation), 0.35 % Bovine Serum Albumin (Sigma) and mannitol to 600 mOsmol. For enzyme removal, the preparation was washed twice with 2 mM CaCl_2_, 2 mM MgCl_2_, 10 mM MES (pH 5.5), and mannitol to 600 mOsmol. For protoplast liberation, the preparation was incubated with 2 mM CaCl_2_, 2 mM MgCl_2_, 10 mM MES (pH 5.5), and mannitol to 280 mOsmol. Filtering the suspension (through a 80 μm nylon mesh) yielded protoplasts. For transient expression, protoplasts were co-transformed as described by Haswell et al. (2008)^17^. Silent protoplasts obtained from quintuple mutant (Δ5) Arabidopsis calli were co-transformed with 2.5 μg 35Sp::GFP in the p327 vector and with 10 μg 35Sp::MSL10 in the pAlligator2 vector. We only used fluorescent protoplasts, indicating a co-transformation, for patch-clamp experiments. As controls for transfection, we tested patches from Δ5 cells transfected with soluble GFP alone (n= 5) and found no mechanically activated currents in the pressure range from 0 to 60 mmHg.

### Electrophysiology

Patch-clamp experiments were performed as described at room temperature with a patch-clamp amplifier (model 200A, Axon Instruments, Foster City) and a Digidata 1322A interface (Axon Instruments). Currents were filtered at 1 kHz, digitized at 4 kHz, and analyzed with pCLAMP8.1 and Clampfit 10 software. During patch-clamp recordings, the membrane potential was clamped at −186 mV and the pressure was applied with a High Speed Pressure-Clamp system^29^ (ALA Scientific Instrument), allowing the application of precise and controlled either pressure pulses or continuous sinusoidal variations in the pipette^11^. Media are designed in order to eliminate stretch-activated K^+^ currents whereas the Ca^2+^ current is negligible compared to that Cl-. Isolated protoplasts were maintained in bathing medium: 50 mM CaCl_2_, 5 mM MgCl_2_, 10 mM MES-Tris, and 0,25 mM LaCl_3_ (pH 5.6). Membrane seal with low resistance (< 1 GΩ) and with unstable current after excision were rejected. The pipettes were filled with 150 mM CsCl, 2 mM MgCl_2_, 5 mM EGTA, 4.2 mM CaCl_2_, and 10 mM Tris-HEPES (pH 7.2), supplemented with 5 mM MgATP. We adjusted the osmolarity with mannitol to 450 mosmol for the bath solution and 460 mosmol for the pipette solution using an osmometer (Type 15, Löser Meβtechnik). Gigaohm resistance seals between pipettes (pipette resistance, 0.8-1.5 MΩ), coated with Sylgard (General Electric) and pulled from capillaries (Kimax-51, Kimble Glass), and the protoplast membranes were obtained with gentle suction leading to the whole-cell configuration, and then excised to an outside-out configuration. The current-pressure relationship data were fitted to a Boltzmann function

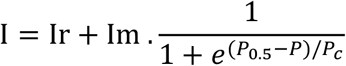

where Ir is the background current at zero pressure, Im is the maximum steady state current intensity, *P*_*c*_ is the slope of the tangent at inflexion point and *P*_0.5_ is the pressure of half activation.

The current activation kinetics were fitted with a mono-exponential function

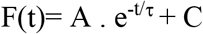

where *A* is current-scale coefficient, τ is the time constant and *C* maximum current intensity.

Mechanosensitive channels respond to the membrane tension, which itself depends on the pipette (and patch) geometry^30,31^. Thus, as the membrane geometry is slightly different from one patch to another (Extended Data Fig. 2a and 2d), Boltzmann functions were determined for each patch individually, prior oscillatory pressure stimulation was applied (Extended Data Fig. 2a). This allows delivering the oscillatory pressure in the same zone of membrane tension sensitivity. The amplitude of the oscillation was +15/-15 mmHg from a mean-pressure baseline that we choose slightly below (5 to 10 mmHg) the P_0.5_ (Fig. 3a).

### Statistical analysis

The data were analysed using Student’s *t*-test and analysis of variance. Comparison of NP(o) at different frequencies (Fig.3 and Extended Fig.4) were analysed with Rank Sum test. NP(o)_osc_ was determined for each oscillatory sinusoidal pressure frequency (0.3 to 30 Hz) and NP(o)_stat_ for each static stimulation prior to frequency stimulation. In Fig. 3b we present the ratio NP(o)_osc_/ NP(o)_stat_ called NP(o) Ratio. The same logic is applied for T_open_ *ratio* and T_close_ ratio Fig. 3c and d.

### Cloning and genetics

All plasmid constructs were made with Gateway technology (Life Technologies). The *MSL10* cDNA was cloned previously into pENTR/D-TOPO^17^. This pENTR construct was then used in recombination reactions with pAlligator^32^ to create the MSL10 protein overexpression construct (*p35S::MSL10*). This construct was used for transient expression in protoplasts obtained from the quintuple mutant *msl4;msl5;msl6; msl9;msl10* (*∆5* + *MSL10*).

### Modeling

We modeled MSL10 as a 2 states channel: an open one (O), in which the channel is activated, and a closed one (C) in which the channel is completely closed. The equilibrium between the 2 states is given by the classical chemical reaction:

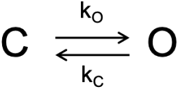

with k_o_ and k_c_ the opening and closing rates, respectively. Opening (resp. closing) rate increases (resp. decreases) exponentially with the applied pressure, describing the mechano-sensitivity in the Arrhenius framework. This model doesn’t contain any specific oscillatory sensitivity, but the reaction rates are affected by the changes in pressure. We then adjusted the four constants of the model to reproduce the experimental data (Fig. 4a).

### Accession numbers

*MSL4*: At1g53470 (SALK_142497, msl4-1)

*MSL5*: At3g14810 (SALK_127784, msl5-2)

*MSL6*: At1g78610 (SALK_06711, msl6-1)

*MSL9*: At5g19520 (SALK_114626, msl9-1)

*MSL10*: At5g12080 (SALK_076254, msl10-1)

## Acknowledgements

This work is supported by the grant ANR-09-BLAN-0245-03 from the Agence Nationale de la Recherche (ANR, project SENZO) and the grant ANR-11-BSV7-010-02 from the Agence Nationale de la Recherche (ANR, project CAROLS). We would like to thank Dr. Sébastien Thomine (I2BC, Gif-sur-Yvette, France) and Dr. Alexis De Angeli (I2BC, Gif-sur-Yvette, France) for their help and for insightful discussions, Dr. Elizabeth Haswell (Washington University, St. Louis, USA) for providing quintuple msl mutant and GUS reporter lines, Gwyneth C. Ingram for corrections on the manuscript (ENS, Lyon, France).

## Author Contributions

D.T performed experiments; T.G generated MSL10 lines; M.G, N.L.F, B.M and E.dL were involved in study design; D.T, J.M.A and J.M.F designed the study; D.T, J.M.A and J.M.F analyzed data; D.T and J.M.F wrote the paper. All authors discussed the results and commented on the manuscript.

## Author information

The authors declare no competing financial interest. Correspondence and requests for materials should be addressed to J.M.F.

## Additional information

**Supplementary Figure 1 | MSL10 pressure dependence of activation time constants**.

**Supplementary Figure 2 | Determination of current-pressure relationship of MSL10 channel**.

**Supplementary Figure 3 | MSL10 channel opening occurred almost exclusively during the upper phase of the stimulation period**.

**Supplementary Figure 4 | Patch clamp recording of MSL10 elicited by oscillatory pressure of 0.1 Hz, 1 Hz, 3 Hz**.

**Supplementary Figure 5 | MSL10 channel characteristics obtained on an individual patch using a protocol alternating oscillatory and static stimulation**.

**Supplementary Movie 1 | Oscillatory movement of an Arabidopsis thaliana plant after elicitation by an air pulse. Slow motion X 16**.

**Supplementary Movie 2 | Activation of the mechanosensitive channel MSL10 elicited by oscillatory pressure of 0.1 Hz, 1 Hz, 3 Hz**.

