## Supplementary Figures for "Transduction of mechanical cellular oscillation by the plasma-membrane mechanosensitive channel MSL10"

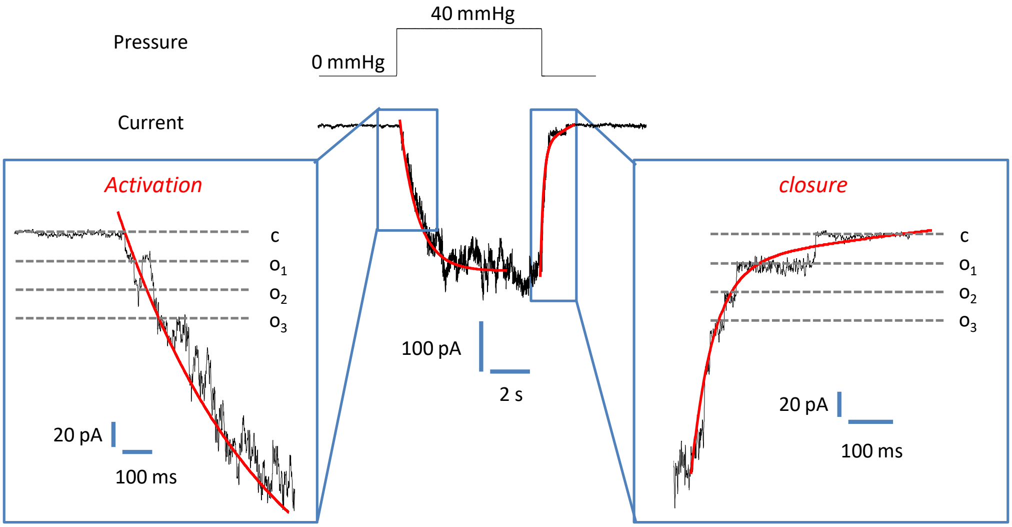


**Supplementary Figure 1** | **MSL10 pressure dependence of activation time constant.** Typical analysis of activation current from quintuple mutant expressing MSL10 elicited by pressure pulses. Current is fitted with a mono-exponential function. Insets show single current transitions happening during pressure steps. Opening (left inset: o1, o2 and o3) and closing (right inset: o3, o2 and o1) of three channels can be seen in this recording.

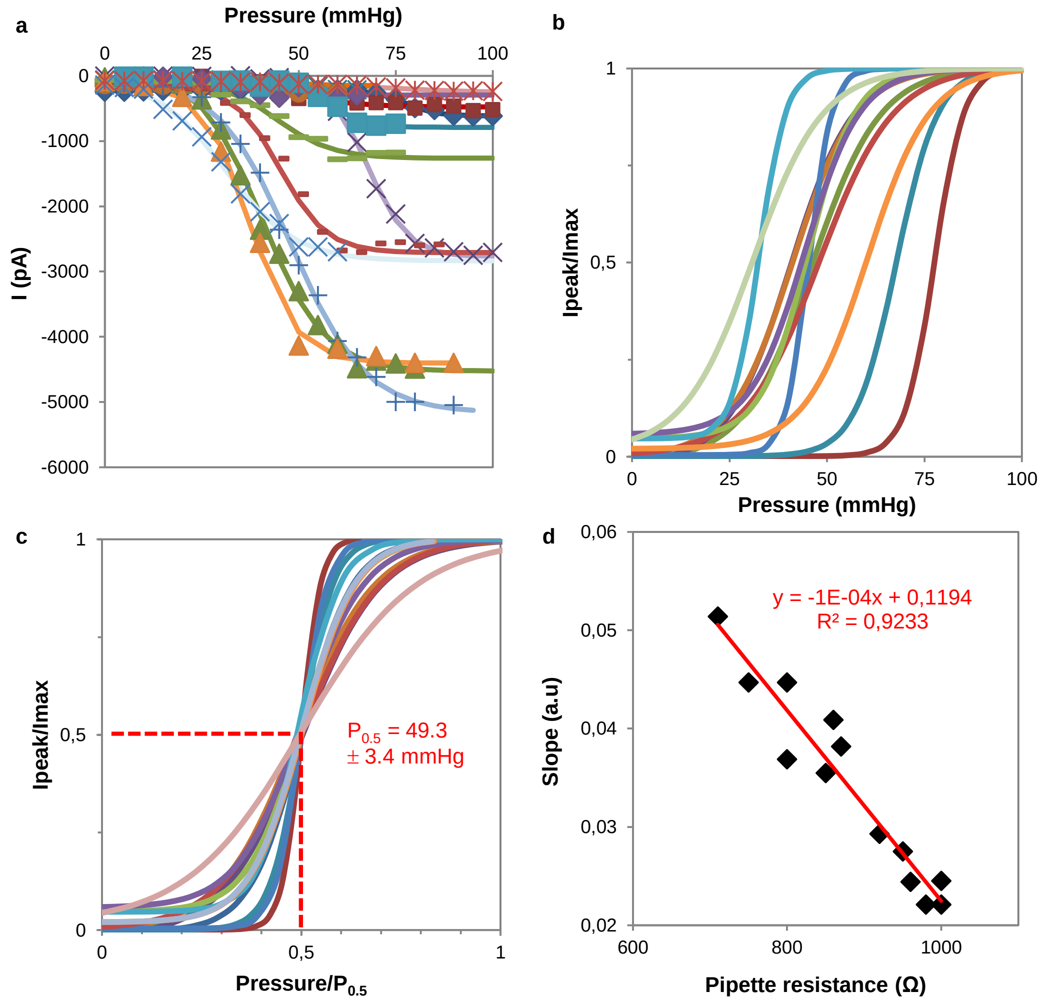


**Supplementary Figure 2** | **Determination of current-pressure relationship of MSL10 channel.**
**a,** Current-pressure relationships recorded in different patches and fitted with a Boltzmann function. Although single activities of MSL10 channels are observed in each recording, the amplitude of the current varies from one patch to another. This reflects different channel densities due to heterogeneity of expression levels between protoplasts. **b,** In order to compare patches, we normalized each curve by maximum current amplitude ($I_{max}$) and **c,** the curves were then normalized to the half-maximal activation point ($P_{0.5}$) from different patches. **d,** Relationship between the slope of the Boltzmann function at $P_{0.5}$ as a function of pipette resistance in our ionic conditions. A linear dependence of pipette resistance (geometry) on Boltzmann curve “steepness” for MSL10 channel activity is observed. This is in accordance with Laplace’s law showing that the MSL10 channel is regulated by membrane tension, which is a general feature for MS channels[33].

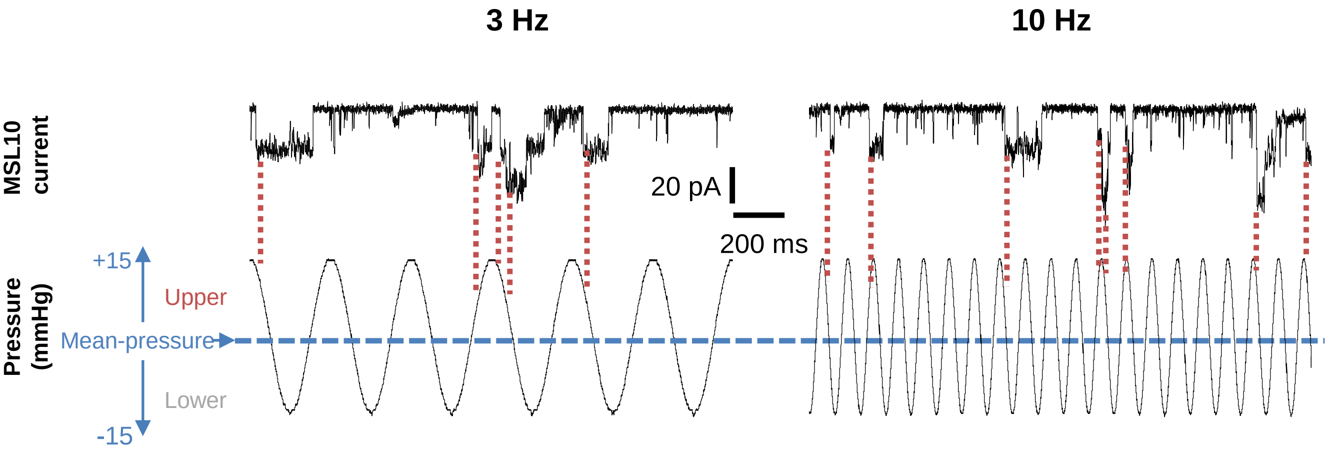


**Supplementary Figure 3** | **MSL10 channel opening occurred almost exclusively during the upper phase of the stimulation period.** Example of recording for stimulation at a frequency higher than 1 Hz (in this recording 3 and 10 Hz) we observed misfiring. Although it is the positive (not negative) phase of the pressure which elicit channel opening, at 3 and 10 Hz not each positive phase is efficient
The membrane potential is clamped at -186 mV. MSL10 protein is transiently expressed in quintuple msl4;msl5;msl6;msl9;msl10 mutant (Δ5) protoplasts. Ionic conditions are described in the Materials and Methods

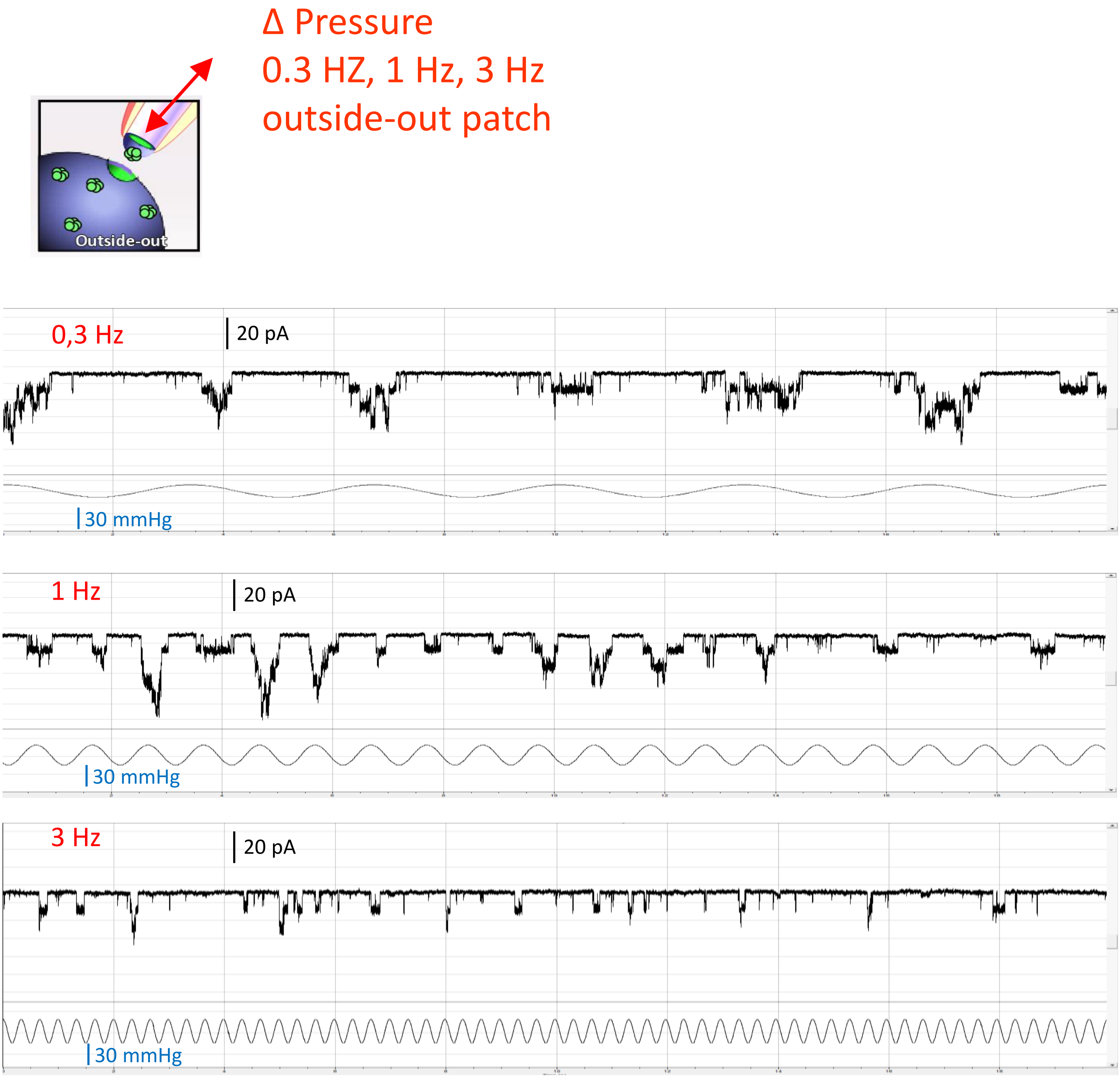


**Supplementary Figure 4** | **Patch clamp recording of MSL10 elicited by oscillatory pressure of 0.3 Hz, 1 Hz, 3 Hz.**

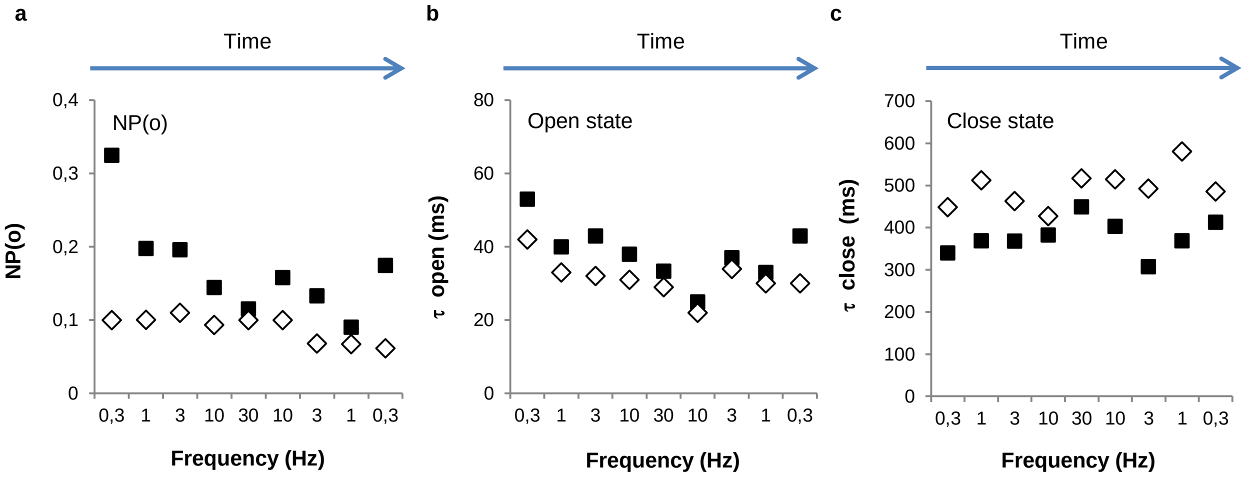


**Supplementary Figure 5** | **MSL10 channel characteristics obtained on an individual patch using a protocol alternating oscillatory and static stimulation.** The same protocol as Figure 3a alternating oscillatory and static stimulation was applied to excised outside-out-patches **a-c,** Representative single patch analysis at different frequencies (*black square*) compared to static stimulation (*white square*). Effects of frequency stimulation on **a,** open probability NP(o), **b,** open state time constant $\tau_{open}$ and **c,** closed state time constant $\tau_{close}$ .

The membrane potential is clamped at -186 mV. MSL10 protein is transiently expressed in quintuple *msl4;msl5;msl6;msl9;msl10* mutant (*Δ5*) protoplasts. Ionic conditions are described in the Materials and Methods.
